# Context-dependent selection and genetic facilitation and constraint on rosette diameter and herbivore resistance across European outdoor common gardens under ambient and reduced precipitation in *Fragaria vesca*

**DOI:** 10.64898/2026.02.12.705624

**Authors:** Ivan M. De-la-Cruz, Carolina Diller, Femke Batsleer, Dries Bonte, Timo Hytönen, José Luis Izquierdo, Sonia Osorio, David Posé, Aurora de la Rosa, Martijn L. Vandegehuchte, Anne Muola, Johan A. Stenberg

## Abstract

The expression of plant defensive traits against herbivores often incurs costs to other essential functions, such as growth and reproduction. Understanding how selection acts on putatively functional traits that are expected to trade off across space and time is therefore critical for predicting evolutionary responses to ongoing and future environmental change. Here, we used multiple replicated genotypes of woodland strawberry (*Fragaria vesca*; Rosaceae) grown over two years in three outdoor common gardens in Spain, Belgium, and Sweden. In each garden, genotypes were exposed to both a reduced-precipitation treatment simulating drought and an ambient precipitation treatment. We estimated directional and correlational selection on rosette diameter (a proxy for growth) and herbivore resistance (measured as the inverse of chewing damage) using fruit and stolon production as proxies for sexual and asexual fitness across all environments (i.e., every site × year × treatment combination). We then combined selection gradients with environment-specific genetic (co)variance among genotypes to quantify the expected response to selection (Δz = *Gβ*) and to identify covariance-driven constraints or facilitation therein. Selection consistently favored larger rosette diameter for both fitness proxies across nearly all environments, which, in combination with genetic covariances among genotypes, resulted in a general evolutionary response toward increased rosette diameter, with the strongest response at the wettest site (Belgium). In contrast, selection on resistance and the corresponding among-genotype evolutionary responses were strongly context-dependent. Correlational selection on rosette diameter × resistance occurred in only a few environments, primarily under reduced precipitation. Environment-dependent genetic covariances constrained or facilitated selection on both traits only at the site with the highest herbivory (Sweden) under drought conditions. Overall, our results reveal a context-dependent interplay between selection and genetic architecture, underlining the difficulty of predicting evolutionary trajectories under environmental change, and highlighting how spatially and temporally variable conditions may maintain standing genetic variation in plant traits.

## Introduction

Plants must often balance their allocation of limited resources between growth and defense against herbivores and pathogens (Coley et al. 1985; Züst and Agrawal 2017; Monson et al. 2022). Growth enhances resource acquisition and competitive ability, whereas defense protects existing tissues through mechanisms such as the production of toxic secondary metabolites that deter, harm or kill herbivores (Züst and Agrawal 2017; Monson et al. 2022). Because nutrients are often limited in nature, investment in defense typically reduces resources available for growth or reproduction, leading to common—though not universal—growth–defense trade-offs due to allocation costs (Coley et al. 1985; Züst and Agrawal 2017; Giolai and Laine 2021; Monson et al. 2022). The trade-offs between growth and defense may also have a genetic basis through pleiotropy or linkage disequilibrium (genetic costs; Züst and Agrawal 2017). Alternatively, simultaneous expression of growth and defense traits may be constrained by environmental conditions (ecological costs; Macel 2011; Züst and Agrawal 2017). For instance, drought or heat stress can suppress plant defenses and alter the cost–benefit balance of plant resistance (Zhao et al. 2025). Thus, trade-offs between defense and growth are often shaped by resource-allocation strategies, genetic correlations, and environmental heterogeneity, which together can generate a mosaic of evolutionary trajectories for these traits across populations (Thompson 2005; He et al. 2022; Monson et al. 2022). However, although growth–defense trade-offs are well documented, comparatively little is known about whether selection gradients align with the underlying genetic (co)variation in these traits, and whether such alignment persists across space, time, and environmental stressors such as drought (but see Züst and Agrawal 2017 and references therein).

Predicting the evolutionary trajectories (response to selection; *Gβ*) of growth and defense requires quantifying selection on each trait and on their combinations, together with the genetic architecture linking them; i.e., evaluating the genetic correlations among defense and growth (Eroukhmanoff and Svensson 2011; Milocco and Salazar-Ciudad 2022; Mallard et al. 2023). These genetic correlations can be detected via the variance–covariance among genotype mean traits (the G matrix) (Agrawal and Stinchcombe 2009; Wise and Rausher 2013; Mallard et al. 2023). Thus, assessing how selection aligns with the G-matrix provides crucial insights into the predicted evolutionary trajectories of growth and defense. For example, even when selection favors both growth and increased defense, a negative genetic covariance (e.g., larger rosettes genetically linked with lower resistance) can counteract these responses and channel evolution along the line of least genetic resistance, thereby slowing down adaptation toward increased growth (i.e., genetic constraint; Coley et al. 1985; Züst and Agrawal 2017). In contrast, covariance aligned with selection can facilitate or increase the multivariate change (Wise and Rausher 2013). There is no universal rule: genetic correlations sometimes constrain adaptation, and other times facilitate it (Wise and Rausher 2013). Moreover, if the G matrix shifts across environments (genotype-by-environment interaction) (Eroukhmanoff and Svensson 2011; Mallard et al. 2023), a tight trade-off in one environment may relax or reverse in another. A genotype’s capacity to respond to selection may thus be itself environment-dependent (Eroukhmanoff and Svensson 2011; Agrawal et al. 2012; Mallard et al. 2023). Nevertheless, few studies have tested whether environment-dependent genetic constraints shape selection gradients on growth and resistance when fitness is partitioned into sexual and asexual components, as in many clonal plants.

In many clonal plants, fitness accrues through both sexual reproduction and asexual propagation, and selection on growth and defense may therefore differ between these components of fitness (Monson et a. 2022). Yet phenotypic selection analyses that explicitly quantify selection via both sexual and asexual fitness proxies remain comparatively scarce, limiting our ability to predict how growth–defense interactions evolve when reproduction occurs through multiple modes (Huot et al. 2014; He et al. 2022; Monson et a. 2022). Here, we used the woodland strawberry, *Fragaria vesca* (Rosaceae) to (1) estimate whether directional and/ or correlational selection acted on plant growth and/or resistance using fruit and stolon production as proxies for sexual and asexual fitness, respectively, (2) test whether selection aligns with the among-genotype genetic variance–covariance for both traits; i.e., the predicted evolutionary response, and (3) evaluate instances in which selection gradients on growth and resistance are constrained or facilitated across different environments, including a reduced-precipitation treatment. To this end, we grew 16 wild *F. vesca* genotypes for two years in three outdoor common gardens located in Spain, Belgium, and Sweden, which vary in environmental conditions such as temperature, precipitation, and herbivory (Figure 1A,B,C,D,E). The woodland strawberry genotypes originate from natural populations spanning the species’ European latitudinal range, providing broad standing genetic variation with which to evaluate the evolutionary response potential in growth and resistance, as well as the underlying genetic synergies or constraints.

**Figure 1.**
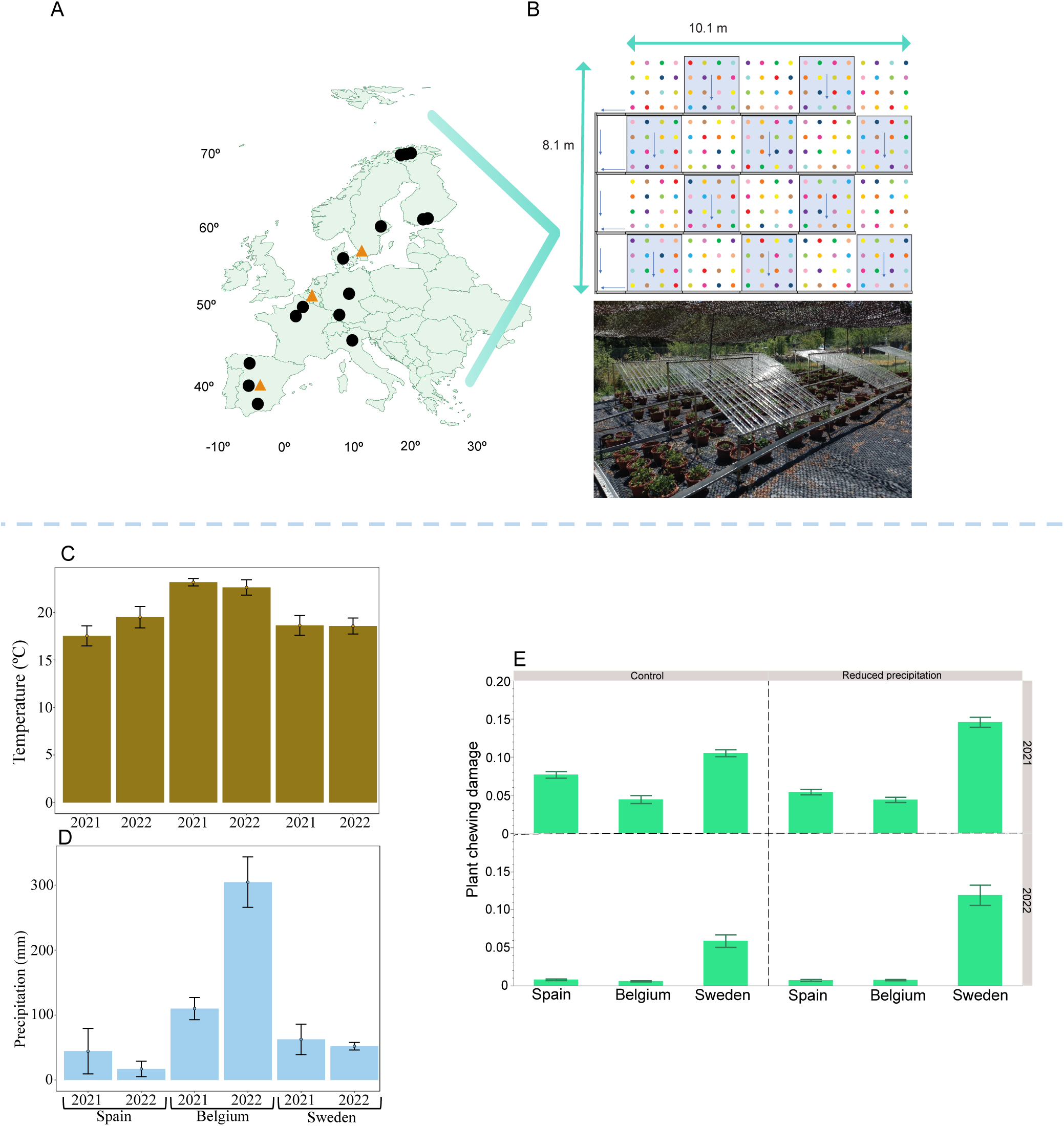
(A) The location of the three experimental sites [orange triangles, from south to mid-and northern Europe: Spain (Rascafría), Belgium (Gontrode), and southern Sweden (Alnarp)] and the origin of each of the 16 genotypes used in the study (black circles; for genotype labels and their latitude and longitude of origin, see Supplementary Table S2). (B) The experimental design for each study site. Each colour represents one of the 16 genotypes. Rainout shelters (blue shaded areas) excluded 50% of the incoming precipitation. The blue arrows indicate the flow direction of the rainwater through the drainage pipes. (C, D) Mean temperature (C) and precipitation (D) and their standard errors in the period April–August at each site (see also Supplementary Table S1). (E) Mean plant chewing damage and its error bars are shown for each site under the control and reduced-precipitation treatments for 2021 and 2022. Credit for picture 1B: Martijn L. Vandegehuchte.

## Materials and methods

The woodland strawberry, *Fragaria vesca*, is a perennial plant species that occurs throughout the Northern Hemisphere in semi-open habitats such as forest clearings and along forest edges (Hancock 1999). It reproduces both sexually via hermaphroditic flowers and asexually (clonally) by forming aboveground stolons (Hancock 1999; Schulze et al. 2012; Muola and Stenberg 2018). The main growing season is between March and August, depending on the geographic location (see Supplementary Table S1).

Three study sites (Fig. 1A) were located along a south–north gradient, spanning the southern and mid-northern portions of the continental European distribution of *F. vesca* (Supplementary Table S1). The study sites were central Spain (Rascafría 40°54′17.941″N, 3°52′46.31″W), Belgium (Gontrode 50°59′0.581″N, 3°47′50.248″E), and southern Sweden (Alnarp 55°38′59.99″N, 13°03′60.00″E). This selection ensured a diverse range of precipitation regimes, temperatures, and herbivore communities across sites (Figure 1C, D, E and Supplementary Table S1). The 16 plant genotypes used in this study originated from distinct wild populations along the continental European south-north gradient (Supplementary Table S2). Complete details on plant growth conditions (including sowing and propagation of plant material), maintenance, and experimental setup are provided in De-la-Cruz et al. (2025a,b). Twenty clonally propagated plantlets per genotype were planted at each site, each in a separate pot, except for one genotype (GER3), for which only 10 replicate plants per site were available.

### Experimental set-up

The plants (*n* = 310 per study site; *n* = 930 in total) were divided into two treatments, reduced precipitation and control, and exposed to the sites’ natural conditions (Figure 1B). At each site, 10 plants from each of 15 of the genotypes, and 5 from genotype GER3, received a reduced precipitation treatment, while the other 10 plants per genotype (5 for genotype GER3) served as controls. A split-plot design was used, with the genotype serving as a split-plot factor (one plant per genotype within a plot; hereafter referred to as a block) and the reduced precipitation treatment applied to whole plots (blocks; Figure 1B). Pots were spaced 50 cm apart in all directions in a regular grid formation (Figure 1B). The reduced precipitation treatment consisted of rainout shelters that reduced the incoming precipitation by 50% (Figure 1B; Yahdjian and Sala 2002). Plants receiving the control treatment were placed in blocks without shelters (Figure 1B). The entire experimental area was covered with MyPex® weed membrane (Don and Low Ltd) to prevent weed growth and fenced with fine-mesh chicken wire (also dug into the soil) to prevent mammalian herbivory. During the summer (June − August), the humidity was measured after 6 − 8 days without rain (or after 3 days without rain if the maximum daily temperatures were +30°C or above) with a soil moisture meter (Fieldscout TDR 150, Spectrum Technologies, Inc.). If the average soil moisture of 20 randomly chosen pots (10 per treatment) was lower than 10% of the volumetric water content (VWC), additional water was supplied: 1 L and 0.5 L to plants under control and reduced precipitation conditions, respectively. Thus, plants in the reduced precipitation treatment always received ∼50% of the amount of water compared to the control plants. Soil moisture was also measured during the growing season (June, July and August) in two different sections of all plant pots during both years. The average soil moisture (%) for the control and the reduced precipitation treatment across all sites was as follows: control^(2021)^ = 25.44 ± 0.52, reduced precipitation^(2021)^ = 11.56 ± 0.40, control^(2022)^ = 28.40 ±0.48 and reduced precipitation^(2022)^ = 12.08 ± 0.37 (see also Supplementary Table S1). These data confirmed that the shelters effectively served as a reduced precipitation treatment (Supplementary Table S1).

### Data collection

#### Leaf chewing damage

The leaf area damaged by chewing herbivores was scored visually for 10 randomly chosen leaves per plant once in June, July and August of 2021 and 2022. We then calculated the mean plant damage across the three months (n = 30). Chewing herbivore damage was scored as proportional leaf area removed and recorded as fractions of a leaflet or, for smaller areas, as decimal values following these criteria: 1/3 of a leaf (i.e., one leaflet), 1/6 of a leaf (i.e., half a leaflet), 1/12 or 1/24 of a leaf (i.e., one-quarter or one-eighth of a leaflet), and specific decimal proportions for smaller areas (0.01, 0.02, or 0.03). To minimize observer bias, participants were trained during bespoke workshops to assess chewing damage on woodland strawberry leaves until estimates converged across participants. In addition, each observer was trained by using the web application https://zaxherbivorytrainer.com to estimate the amount of leaf damage on randomized images of leaves. Estimates within 0 − 5% of the actual answer were considered accurate. See Figure 1E for the mean leaf damage values per site across treatments and years.

#### Rosette diameter and reproductive traits

For each plant, rosette diameter was used as a proxy for vegetative growth and measured horizontally with a ruler (cm). Proxies of sexual and asexual fitness were estimated as the number of ripe fruits and stolons (Muola and Stenberg 2017; De-la-Cruz et al. 2025a), respectively. Rosette diameter and counts of fruits (including those fallen into the soil pots) and stolons were recorded monthly in June, July, and August of each study year (2021 and 2022). Only fruits without visible herbivore damage or pathogen infection were included, as such damage can reduce seed number and quality and thereby bias fitness estimates. Fruits and stolons were systematically removed after counting to prevent double counting and, in the case of stolons, to avoid growth into neighboring pots. Fruit removal is unlikely to induce compensatory responses (Hilty et al. 2021), as ripe fruits naturally detach from the plant over time. Rosette diameter was averaged across the three sampling times within each year for downstream analyses (see below). Fruit and stolon counts were summed across the three sampling times within each year and used as annual proxies of plant fitness for subsequent analyses (see below).

### Statistical analysis

#### Phenotypic selection on resistance and rosette diameter across environments

All analyses were conducted in R v4.4.0 (R Core Team 2021). We used the inverse of plant damage as a proxy for plant resistance (calculated using the operational definition 1 – mean proportion of total leaf area damaged by chewing herbivores; Simms and Rausher 1987; Núñez-Farfán and Dirzo 1994). This operational resistance measurement reflects only realized chewing damage (Simms and Rausher 1987; Núñez-Farfán and Dirzo 1994). However, because herbivores often preferentially attack larger, healthier plants that offer greater biomass and nutrient availability (i.e., increased apparency; Price 1991; Hella Schlinkert et al. 2015), and because plant resistance may align with a latitudinal cline (Woods et al. 2012), we adjusted our measure of plant resistance by using a generalized linear mixed model (GLMM) to obtain size-and latitude-independent estimates of herbivore resistance within each environment (i.e., each site × treatment × year combination). To this end, for each environment, we fitted a GLMM (beta error distribution with a logit link), using proportional plant damage as the response variable and rosette diameter and latitude as fixed effects or predictors. Genotype and block were included as random effects to account for genetic and spatial replication. From each model, we extracted environment-specific slopes for rosette diameter (*b*_diameter_) and latitude (*b*_latitude_), quantifying the effects of plant size and geographic origin on herbivore damage. We then calculated individual-level adjusted damage by subtracting the predicted contributions of rosette diameter and latitude from the observed logit-transformed damage: Adjusted damage = logit(Damage_observed_) − (*b*_diameter_ × rosette diameter) − (*b*_latitude_ × latitude). This value represents the expected damage a plant would experience if it had the mean rosette diameter and latitude of its origin (Stinchcombe et al. 2008). Adjusted damage was back-transformed using the inverse logit function, and adjusted resistance was defined as 1 − adjusted damage. This metric enabled selection analyses on herbivore resistance that are unbiased by predictable variation in plant apparency and geographic origin (Price 1991; Hella Schlinkert et al. 2015). Likewise, to remove potential geographic clines in growth (as observed by De-la-Cruz et al. 2025a in the same dataset analyzed here), rosette diameter was residualized against latitude within each environment using a mixed-effects model with genotype and block included as random effects, yielding a latitude-independent estimate of plant size (Woods et al. 2012).

We then performed proxies of directional and correlational phenotypic selection analyses on resistance and rosette diameter (adjusted; see above) using two different GLMMs, one for each fitness measures (i.e. for fruit and stolon production). In each model, the response variable was fruit or stolon production. To estimate selection on *relative* fitness proxies within each environment (i.e., every site *×* year *×* treatment combination), we computed the environment-specific mean for fruit and stolon production and used its natural logarithm as an offset in the two models. The GLMMs were fit using a negative binomial distribution with a log link (variance function “nbinom2”) using glmmTMB v1.1.13 (Brooks et al. 2017; McGillycuddy et al. 2025). The structure of the models was specified as follows:

Fruits/stolons = Offset (Log (Mean fruits/stolons)) + (resistance + rosette diameter + resistance × rosette diameter) * site * year * treatment.

Resistance and rosette diameter values were standardized to mean 0 and standard deviation of 1 within each environment (i.e., site × treatment × year combination). In these models the independent effect (or additive effect) of resistance or rosette corresponds to directional selection gradient proxies, while the interaction (synergistic/antagonistic effects) between resistance and rosette corresponds to the correlational selection proxies. We included the block, genotype × site, genotype × year, genotype × treatment, genotype × site × year, genotype × year × treatment, genotype × site × treatment, and genotype × site × year × treatment as random effects. Incorporating the random effects into these GLMMs allowed each genotype to have its own baseline shift in fitness in each environment.

Because the model uses a log link and an offset, coefficients on the latent (log) scale correspond to multiplicative effects on expected fruit or stolon production relative to the environment mean (McCullagh and Nelder 1989; de Villemereuil et al. 2016). Thus, we used emtrends v1.3.5.1 to obtain, for each environment, the marginal slopes (“trends”) for resistance, rosette diameter, and their interaction on the latent scale, along with standard errors and Wald tests. We back-transformed the latent-scale gradients to the raw scale by using: *βraw* = *e*^!^ − 1 (McCullagh and Nelder 1989). Standard errors and 95% confidence intervals were also calculated for the raw scale selection gradients. The generalized linear coefficients (i.e., the selection gradients; *βi*, Lande and Arnold 1983) obtained from the models represent the magnitude and direction of selection acting on resistance, rosette, resistance × rosette in comparable units (standard deviations) across years, sites, and treatments (Lande and Arnold 1983; Wise and Rausher 2013).

#### Among-genotype variance–covariance matrix per environment

For each environment (every site × year × treatment combination), we performed a two-response bivariate bayesian GLMM in the R package brms v2.23.0 (Bürkner 2017) with the cmdstanr v0.9.0 interface (Gabry et al. 2025). Resistance was modeled with a Beta distribution and logit link as follows: Resistance ∼ Block + (1 | Genotype). Rosette diameter was modeled assuming a Gaussian distribution and identity link as follows: Rosette diameter ∼ Block + (1 | Genotype). Block was included as a fixed effect for both traits to control for micro-environmental heterogeneity/structure variance within sites. Genotype as random effect for the multivariate GLMM allowed the model to estimate the among-genotype covariance between traits. Residual cross-trait correlation was disabled, ensuring that cross-trait association is attributed to the genotype effects rather than to residual error (de Villemereuil et al. 2016). For each environment-specific model, we ran four chains, with 8,000 iterations per chain, warm-up to 25% of the total, adapt_delta = 0.95 and max_treedepth = 15 and fixed random seed of 1001. For each fitted environment-specific model, posterior draws (each draw is one possible 2 × 2 variance-covariance matrix from the MCMC bayesian GLMM; ∼288,000 draws in total across all environments) were converted to a latent-scale G-matrix (G_lat_); a 2 × 2 genotype-level variance–covariance matrix G on the latent scale (logit scale for resistance; identity for rosette diameter). We converted the latent covariance and variances matrices (G_lat_) to observed scale (i.e., the natural measurement scale of each trait; G_obs_) using *Qgmvparams* function of QGglmm v0.8.0 R package (de Villemereuil et al. 2016). Thus, we obtained an observed-scale genetic (co)variance matrices (G_obs_) per environment, consistent with the link functions and distributional assumptions of the two traits (de Villemereuil et al. 2016).

The observed G matrices (G_obs_) were further standardized to enable comparisons across environments. For each environment, we computed the phenotypic standard deviation vector and formed a diagonal scaling matrix (Khachiyan and Kalantari 1990). The observed-scale matrices (G_obs_) were then standardized as G_st_ = D^−1^G_obs_D^−1^ (Khachiyan and Kalantari 1990), where D^−1^ is the diagonal scaling matrix for resistance and rosette diameter for each environment and G_obs_ is the observed G matrix for that environment (see above). The G_st_ matrices were summarized by calculating the mean among-genotype variance-covariance matrix (G_st-mean_) of resistance and rosette diameter together with their 95% credible intervals for each environment. Diagonal elements quantify genetic variance for each trait; the off-diagonal element reflects the sign and magnitude of the genetic covariance between resistance and rosette.

#### Evolutionary response potential

We estimated the among-genotype evolutionary response potential for each environment using Lande’s equation, Δz = *Gβ* (Lande 1979; Milocco and Salazar-Ciudad 1999), where *G* is the 2 × 2 environment-specific among-genotype variance–covariance matrix (G_st_; see above) and *β* is the vector of directional selection gradients on resistance and rosette size for that environment. From the Lande’s equation, we retained both components: Δresistance and Δrosette diameter (the among genotype evolutionary response for resistance and rosette diameter for each environment). Δresistance and Δrosette diameter are expressed in phenotypic standard deviation units. We stress that we interpret Δz = *Gβ* as the among-genotype evolutionary response potential in every environment rather than population-specific evolutionary predictions (Milocco and Salazar-Ciudad 1999). In other words, Δz describes the expected change in the trait mean if selection acted on differences among the genotypes studied within a given environment (every site × treatment × year combination). These values are not forecasts of population-level evolutionary trajectories, as they do not account for genotype/allele frequencies or other population-specific demographic processes (Milocco and Salazar-Ciudad 1999). Positive values indicate a comparative (among-genotype) predicted increase in resistance or rosette diameter mean, while negative values indicate a decrease.

#### Facilitation and genetic constraints

We estimated facilitation and genetic constraints following the approach of Agrawal and Stinchcombe (2009) and Wise and Rausher (2013). From the bayesian GLMMs, we extracted the posterior draws (posterior G_st_ matrices) for each environment (see above). We obtained the directional selection gradients (*β*) for resistance and rosette diameter per environment from the negative binomial GLMM analyses that we described above. For each posterior draw, we again calculated the Δz = *G* × *β* under two scenarios: 1) Full G matrix or Response Vector₁: RV₁ = *G* × *β*, and 2) Diagonalised G matrix (G₀, covariances set to zero) or Response Vector_0_: RV_0_ = diag(*G*) × *β* (Wise and Rausher 2013). Thus, RV_1_ estimates the among-genotype evolutionary response for resistance and rosette diameter considering the genetic covariances among them for each posterior draw, whereas RV_0_ estimates what the evolutionary response would have been if there had been no genetic covariances among them (Smith and Rausher 2008; Agrawal and Stinchcombe 2009). We then quantified the extent to which the predicted response of resistance or rosette size changed when genetic covariances were included (Wise and Rausher 2013). To do so, we compared the predicted change under the full G scenario with that obtained under the no-covariance scenario, as follows: Trait difference (TDiff) = RV₁ trait − RV₀ trait (Wise and Rausher 2013). We then interpreted the results as “facilitation” or “constraint” based on whether covariances increased or decreased the evolutionary response in the direction favored by selection, rather than relying solely on the raw sign of TDiff. We therefore defined a direction-aware covariance effect for each trait by projecting the TDiff onto the corresponding selection gradient, as follows: Facilitation/Constraint (FC) = TDiff × *β*. Under this definition, genetic covariances were classified as facilitating evolutionary change for resistance or rosette diameter when FC > 0, indicating that covariances increase the magnitude of the predicted response in the direction of selection (Agrawal and Stinchcombe 2009; Wise and Rausher 2013). Conversely, covariances were classified as constraining when FC < 0, indicating that covariances reduce/constraint the predicted response in the direction of selection (Agrawal and Stinchcombe 2009; Wise and Rausher 2013). In other words, we asked if the predicted phenotypic selection was aligned or misaligned with the structure (e.g., antagonistic or synergistic covariance between traits) of the genetic matrix (Wise and Rausher 2013). For each environment, we computed the proportion of draws where each trait experienced facilitation or constraint. An effect was considered significant when ≥ 90% of the posterior draws supported either facilitation or constraint (Lambert 2018).

## Results

### Phenotypic selection on resistance

Significant phenotypic selection on resistance was detected only in Sweden and Spain. In Sweden, under control conditions in 2021, plants with higher resistance produced more fruits (Fig. 2A, Supplementary Table S3). In contrast, in Sweden in 2022 under reduced precipitation, plants with lower resistance had higher fruit production (Figure 2A, Supplementary Table S3). We also found that plants with higher resistance consistently produced fewer stolons in Sweden across both years and treatments and in Spain (only in 2022 under reduced precipitation) (Figure 2B, Supplementary Table S3).

**Figure 2.**
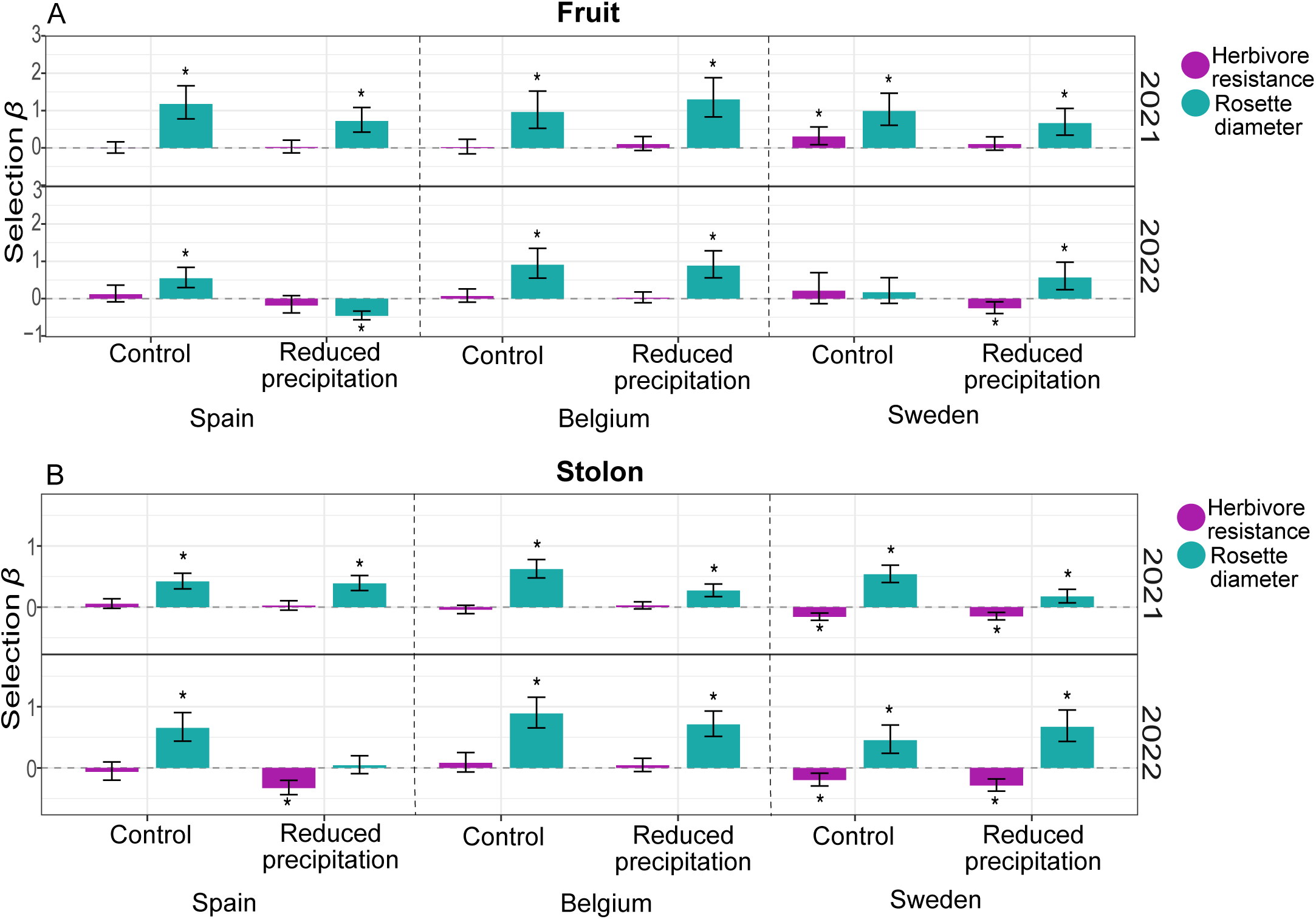
(A) Selection gradients (*β*) for fruit production across the study sites, Spain, Belgium, and Sweden, under control and reduced-precipitation treatments in 2021 and 2022 (site × treatment × year combinations). Purple bars denote selection on chewing herbivore resistance, and teal bars denote selection on rosette diameter. 95% confidence intervals indicate uncertainty around the selection gradients. Asterisks indicate statistical significance (*p* ≤ 0.05). (B) Selection gradients (*β*) for stolon production, with colors and annotations as in panel A.

### Phenotypic selection on rosette diameter

For fruit production, gradients of selection on rosette diameter were positive in 11 of the 12 environments (i.e., in almost every site × treatment × year combination; Figure 2AB, Supplementary Table S3). This pattern suggests that plants with larger rosette diameter had more fruits across sites, treatments, and years (Figure 2A, Supplementary Table S3). The exception occurred in Spain in 2022 under reduced-precipitation conditions, where plants with higher fruit production had smaller rosette diameter (Figure 2A, Supplementary Table S3). In terms of stolon production, selection on rosette diameter showed positive gradients in nearly all environments (almost all significant too), indicating that larger plants had more stolons across sites, treatments, and years (Figure 2B, Supplementary Table S3).

### Among-genotype evolutionary response potential for resistance

Among-genotype evolutionary response for resistance, using fruit production as the fitness proxy, showed a predicted response for higher resistance in Sweden in 2021 under reduced precipitation (Figure 3A). However, in 2022, a predicted response for lower resistance was detected in Sweden under reduced precipitation (Figure 3A), and the same pattern was observed when stolon production was used as a fitness proxy at the same site, year, and treatment (Figure 3B).

**Figure 3.**
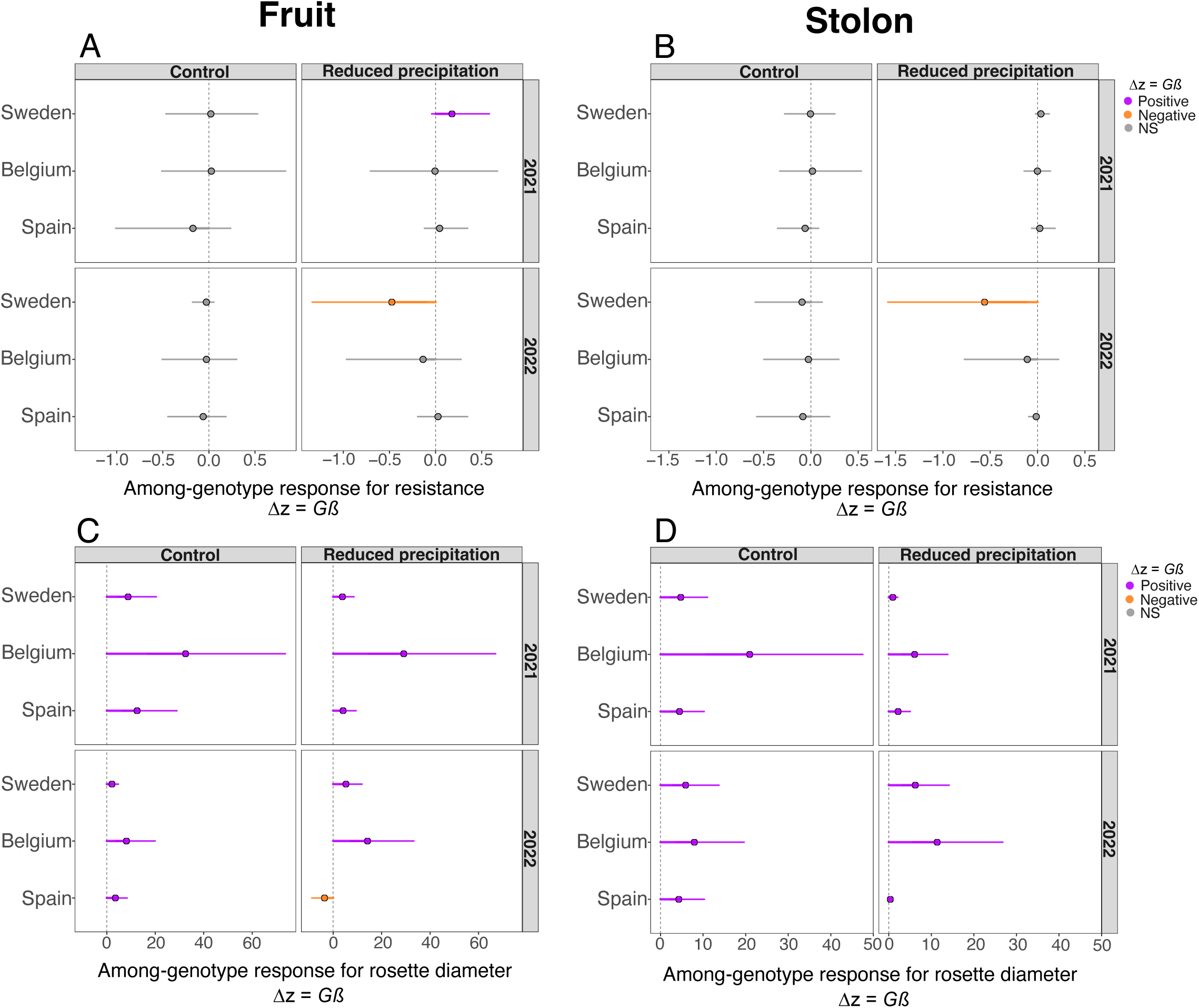
Panels (A–D) show means and 95% confidence intervals (CIs) of the among-genotype predicted evolutionary response to selection (Δz = *Gβ*) under control and reduced-precipitation treatments in 2021 and 2022 across the three study sites: Spain, Belgium, and Sweden. (A) Predicted evolutionary responses (Δz) of herbivore resistance based on selection via fruit production. (B) Predicted evolutionary responses (Δz) of herbivore resistance based on selection via stolon production. (C) Predicted evolutionary responses (Δz) of rosette diameter based on selection via fruit production. (D) Predicted evolutionary responses (Δz) of rosette diameter based on selection via stolon production. Colors indicate whether each mean Δz (±95% CI) is positive (purple), negative (orange), or non-significant (grey). The vertical dashed line denotes Δz = 0.

### Among-genotype evolutionary response potential for rosette diameter

We detected a consistent among-genotype evolutionary response favoring larger rosette diameter across all sites, treatments, and years, using both fitness proxies (Figure 3C,D). The strongest evolutionary response for rosette diameter (both fitness measures) was observed in Belgium across both treatments and years (Figure 3C,D). The exception was found in Spain in 2022 under reduced precipitation using fruit production as fitness measure, where a decrease in rosette diameter was predicted (Figure 3C,D).

### Selection on resistance × rosette diameter interaction

The GLMM for fruit production revealed that in Sweden in 2022 under reduced-precipitation, plants with higher resistance and larger rosettes produced more fruits (Supplementary Figure S1A, Supplementary Table S3). In contrast, in Spain in 2022 under reduced precipitation, plants with lower resistance and smaller rosettes had higher fruit production (Supplementary Figure S1A, Supplementary Table S3). In Belgium, under reduced precipitation, plants with larger rosette diameter and lower resistance produced more fruits in 2021, and the same pattern was observed in 2022, although only marginally significant (Supplementary Figure S1A, Supplementary Table S3). Significant correlational selection on stolon production was only detected in Spain in 2022 under reduced precipitation, indicating that plants exhibiting both higher resistance and larger rosette diameter produced more stolons (Supplementary Figure S1B, Supplementary Table S3).

### Genetic facilitations and constraints

Using fruit production as fitness proxy, we found that in Sweden under reduced precipitation in 2021, the genetic covariances facilitated the selection for an increased resistance and rosette diameter (Figure 4A, Supplementary Table S4). However, in 2022, in Sweden under reduced precipitation, the covariances facilitated selection for reduced resistance (i.e., intensified selection against resistance) and also facilitated selection for larger rosettes (Figure 4A, Supplementary Table S4).

**Figure 4.**
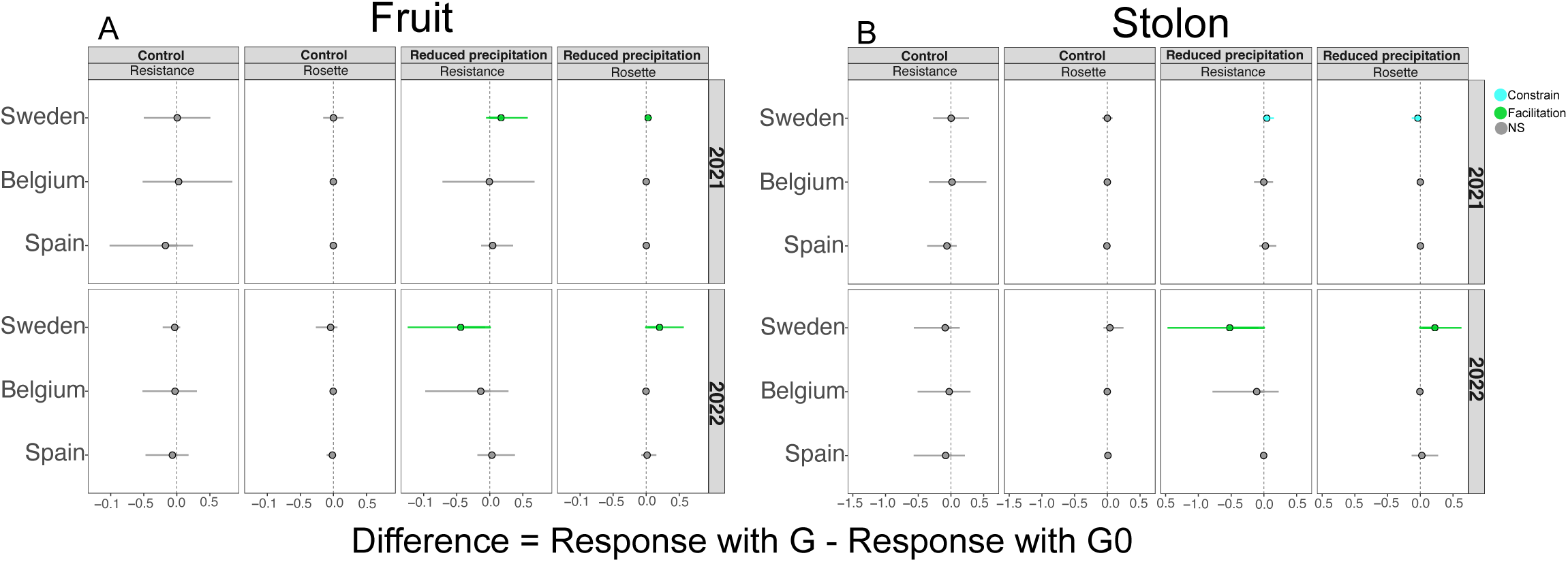
Panels (A) and (B) illustrate per-trait genetic constraints and facilitation, quantified as the difference between predicted evolutionary responses calculated using the full genetic variance–covariance matrix (G) and those calculated with covariances removed (G0). Thus, Difference = Response with G – Response with G0 (see also methods). (A) Median differences and 95% confidence intervals (CIs) based on selection via fruit production across precipitation treatments (control *vs*. reduced precipitation) for each focal trait (herbivore resistance and rosette diameter) in 2021 and 2022 across Spain, Belgium, and Sweden. (B) Median differences and 95% CIs based on selection via stolon production across precipitation treatments and focal traits for 2021 and 2022 across the same sites. The vertical dashed line indicates no effect of genetic covariances (Difference = 0). Colors indicate whether covariances constrain the selection response (blue), facilitate the selection response (green), or have no detectable effect (grey). See also methods and Supplementary Table S4.

Using stolon production as a fitness proxy, genetic covariances in 2021 in Sweden under reduced precipitation constrained selection favoring resistance and also constrained selection for reduced rosette diameter (i.e., slowed selection for smaller rosettes; Figure 4B, Supplementary Table S4). In contrast, in 2022 in Sweden under reduced precipitation, covariances facilitated selection for reduced resistance and facilitated selection for larger rosette diameter (Figure 4B).

## Discussion

This study reveals how environmental heterogeneity interacts with genetic architecture to shape among-genotype evolutionary trajectories of resistance and growth across different environments spanning much of the European range of *F. vesca*.

Across nearly all conditions examined (year × site × treatment combinations), plants with larger rosette diameters consistently showed higher fruit and stolon production. Because genotypes differed in rosette diameter, this pattern implies that, if heritable, mean rosette size would be expected to increase. The strongest predicted among-genotype evolutionary response for larger rosettes occurred in Belgium, likely due to consistently high precipitation across both years, which may have increased resource availability for growth and strengthened selection for larger plants (Hilty et al. 2021). Larger rosette diameters may confer higher fitness through greater photosynthetic capacity, improved resource acquisition, and enhanced competitive ability (Poorter et al. 2012; Ainsworth and Bush 2011; Hilty et al. 2021). An exception was found in Spain under the reduced-precipitation treatment in 2022, where smaller rosettes had higher fruit production. Spain was the driest site in 2022, hence plants may have experienced a more severe drought stress at that site. Under intense water limitation, large rosettes may incur high metabolic costs that reduce fruit production (Eziz et al. 2017). Under such conditions, reduced growth can function as a strategy to limit transpiration and metabolic demand (Eziz et al. 2017; Takahashi et al. 2020).

In contrast to the consistent selection for larger rosette diameter, selection on herbivore resistance was absent or non-significant when either stolon or fruit production was used as a fitness proxy. Likewise, the predicted among-genotype evolutionary response for resistance generally resulted in a non-significant change, except in Sweden under reduced precipitation in 2021, where resistance was predicted to increase. Plants in Sweden experienced higher chewing-herbivore damage than at the other sites in both years and treatments. Under control conditions in 2021 in Sweden, higher resistance was associated with increased fruit production, and we detected a marginally significant positive selection gradient under reduced precipitation, suggesting that a higher resistance conferred higher fitness when herbivore pressure was elevated in 2021. However, in 2022, selection on resistance was not significant under control conditions, whereas under reduced precipitation, higher resistance was associated with lower fruit production. We also detected a predicted evolutionary response against resistance (a predicted shift toward lower resistance) in Sweden in 2022 under reduced precipitation using both fitness proxies. It is possible that the metabolic costs of maintaining defensive traits could outweigh their protective benefits when water/resources are limited (Coley et al. 1985; Züst and Agrawal 2017; Monson et al. 2021). In other words, producing defenses under reduced precipitation when herbivory was high appeared to impose fitness costs. Moreover, we hypothesize that increasing plant size, under both treatments, provided greater benefits for fruit and stolon production than allocating in defense at the herbivory levels observed in Sweden. At these damage levels (∼10% leaf loss), resistance may be more costly than beneficial, particularly under reduced precipitation, and plants may instead cope with herbivory through compensatory growth or tolerance (Nuñez-Farfán et al. 2007; Muola et al. 2010; Züst and Agrawal 2017; Blumenthal et al. 2020), a possibility that needs further study.

Phenotypic selection analyses using stolon production as the fitness proxy further showed that plants with higher resistance produced fewer stolons especially in Sweden across all treatments and years and in Spain in 2022 under reduced precipitation. Using stolon production as a fitness measure adds insights into how resistance was selected against and stresses the importance of allocating resources to growth and clonal propagation rather than to resistance traits. Our findings appear to align with the resource-allocation theory, which suggests that selection for rapid reproduction and larger size can favor fast-growing but weakly defended genotypes (Coley et al. 1985; Monson et al. 2021).

Correlational selection on resistance × rosette diameter (for both fitness proxies) was largely absent in all environments, with four exceptions. In Sweden in 2022 under reduced precipitation, plants with both higher resistance and larger rosettes produced more fruits, suggesting a possible synergistic benefit of investing in growth and defense when water is limited, and herbivory is high (Li et al. 2024). There is empirical evidence from other study systems that under intense herbivory, plants with both higher growth and higher resistance show higher fitness (Carmona and Fornoni 2013; De-la-Cruz et al. 2023). However, it is important to recall that we also found directional selection for reduced resistance in this environment (see above). Thus, we hypothesize that resistance appears beneficial only when accompanied by an allocation for a larger rosette diameter. In other words, we hypothesize that increasing resistance alone is costly—especially under water limitation—unless plants can also allocate resources to growth (Carmona and Fornoni 2013).

In Spain in 2022 under reduced precipitation, correlational selection pointed in opposite directions depending on the fitness proxy. When fruit production was used, the favored trait combination consisted of smaller rosettes and lower resistance; when stolon production was used, the favored combination consisted of larger rosettes and higher resistance. In other words, plants investing more in growth and defense appeared less successful sexually but more successful clonally. These results suggest a trade-off between reproductive modes at the site where the most severe drought and high temperatures were observed. It is possible that larger, better-defended plants allocated all their resources to stolon production, a strategy that may enhance persistence under extreme conditions (Vallejo-Marín et al. 2010; Orive et al. 2017; Blumenthal et al. 2020). In Belgium under reduced precipitation, plants with larger rosettes and lower resistance produced more fruits in 2021, with a similar but marginally significant pattern in 2022. These results from Belgium reinforce that reduced precipitation can promote selective trade-offs (see above). In summary, correlational selection on growth and resistance is not fixed but shaped by reproductive modes and/or ecological context, including spatial variation in herbivory and precipitation.

Genetic correlations can either enhance or constrain the evolutionary adaptive potential, with positive correlations among traits under positive selection accelerating adaptation and negative correlations inhibiting it (Agrawal and Stinchcombe 2009). We detected both genetic constraints and facilitation only in Sweden, and the direction and magnitude of these effects were strongly dependent on environmental context. For fruit production, in Sweden under reduced precipitation in 2021, genetic covariances facilitated the selection for increased resistance and larger rosette diameter, indicating that the G-matrix (e.g., via pleiotropy, linkage disequilibrium; Eroukhmanoff and Svensson 2011; Svensson 2023) may boost their joint adaptive evolution. When stolon production was used as the fitness proxy in Sweden in 2021 under reduced precipitation, genetic covariances constrained evolution for resistance and toward smaller rosettes. Larger rosettes may enable greater allocation to stolon production than smaller rosettes (Orive et al. 2017), and clonality can distribute risk among ramets, promote division of labor, and enhance persistence in stressful environments (Vallejo-Marín et al. 2010; Orive et al. 2017). Thus, plants may have allocated more resources to vegetative growth (thereby reducing evolutionary change toward smaller rosettes), which in the first year could have increased stolon production to persist under drier conditions and herbivory. At the same site in 2022 under reduced precipitation, we found that selection would tend to decrease resistance (facilitation for a reduced resistance), while simultaneously facilitating an increase in rosette diameter. This temporal reversal in the evolutionary responses of resistance highlights how the evolutionary consequences of genetic architecture can shift rapidly with changing selection regimes (genotype-by-environment interactions; Eroukhmanoff and Svensson 2011). These results also show that the reproductive mode plays an important role in shaping the evolution of these traits and highlights the complexity plants face when coping with multiple stressors while allocating resources to both reproductive modes, growth, and resistance (Züst and Agrawal 2017).

In summary, the genetic constraints and facilitations identified only at the site with the highest herbivory (Sweden), together with the context-dependent nature of selection, highlight the difficulty of predicting among-genotype potential responses to climate change without integrating both environmental variation (biotic and abiotic factors) and the genetic architecture.

## Acknowledgements

We thank Juan Antonio Vielva Juez, Director of the Centro de Investigación, Seguimiento y Evaluación del Parque Nacional de la Sierra de Guadarrama, Madrid, Spain for hosting the southernmost (Rascafría/Spain) study site and for logistic support during the experiment. We thank Hans Matheve for helping with fieldwork and data digitalization and Sanne de Jong, Matilda Jützeler and several other assistant researchers who helped us to collect the data during fieldwork. We thanks to UPPMAX (Uppsala Multidisciplinary Center for Advanced Computational Science) of Uppsala University, for providing us access to their computational resources for the Bayesian analyses.

## Author contributions

The study was conceived by IMDC, MLV, AM, JAS, DB, TH, SO, DP. The experiments were performed by IMDC, CD, FB, MLV, ADL, JLI, AM and JAS. The data were analyzed by IMDC. The draft manuscript was written by IMDC with considerable input and revisions from all the authors.

## Open Research

Data supporting this manuscript, and all the scripts/workflow will be deposited in Figshare or Zenodo upon manuscript acceptance.

## SUPPLEMENTARY MATERIALS

**Supplementary Table S1.** Main abiotic environmental conditions occurring at the three experimental sites. Average soil moisture (% of humidity) measured with a FieldScout TDR 150, Spectrum Technologies, Inc. Soil moisture measurements were taken from two opposite sides on the surface of each plant’s pot at two different time points during the growing season of *F. vesca* at each site in 20211 and 2022. Mean annual temperature and precipitation values were retrieved from the closest weather stations: Spain: Agencia Estatal de Meteorología; Belgium: The Royal Meteorological Institute of Belgium; Sweden: Swedish Meteorological and Hydrological Institute. MASL = Meters above sea level. See also Figure 1A-D.

| Site | Year | Mean soil moisture<br>(% of humidity)<br>during growing season |  | Mean<br>temperature<br>(°C)<br>during<br>growing<br>season | Mean<br>rainfall<br>(mm)<br>during<br>growing<br>season | Length of<br>growing<br>season | Flowering<br>period | Altitude<br>MASL |
| --- | --- | --- | --- | --- | --- | --- | --- | --- |
|  |  | Control | Reduced<br>precipitation |  |  |  |  |  |
| Spain | 2021 | 14.23 | 4.97 | 17.53 | 44.26 | April–<br>August | Late<br>April–July | 1200 |
|  | 2022 | 27.57 | 18.81 | 19.49 | 17.06 |  |  |  |
| Belgium | 2021 | 42.86 | 24.52 | 23.18 | 110 | Late<br>March–<br>August | Late<br>April–July | 23 |
|  | 2022 | 38.46 | 12.89 | 22.63 | 304.6 |  |  |  |
| Sweden | 2021 | 16.85 | 5.79 | 18.63 | 62.60 | Late<br>March–<br>August | Late<br>April–July | 9 |
|  | 2022 | 12.83 | 3.03 | 18.56 | 52.03 |  |  |  |

**Supplementary Table S2.** Genotype labels and latitude and longitude of origin. Order is from lowest to highest latitudes.

| Genotype | Latitude | Longitude |
| --- | --- | --- |
| ES2 | 37.7796 | -3.7849 |
| ES20 | 40.2938 | -5.0091 |
| ES13 | 43.1344 | -4.888 |
| IT3 | 45.94 | 10.81 |
| GER100 | 47.96666667 | 7.83333333 |
| FR2 | 48.801407 | 2.130122 |
| FR3 | 50.015654 | 2.6973567 |
| GER3 | 51.4192 | 11.128 |
| LIT3 | 54.5729 | 24.6722 |
| DK1 | 55.5703 | 9.7466 |
| SE6 | 60.1018 | 18.3433333 |
| FIN13 | 60.2292 | 25.0233 |
| FIN39 | 60.4051 | 25.1562 |
| NOR21 | 70.1671 | 24.7561 |
| NOR19 | 70.0301 | 22.0653 |
| NOR8 | 70.0324 | 23.4012 |

**Supplementary Table S3.** Directional and correlational selection gradients on herbivore resistance and rosette diameter across environments (every site × year × treatment combination). The linear selection gradients (*β*) for resistance and rosette diameter and the correlational selection coefficient (Resistance × Rosette) were estimated separately for fruit production (sexual fitness proxy) and stolon production (asexual fitness proxy). Standard errors (SE) and *p*-values testing the null hypothesis of no selection (coefficient = 0) are also shown. Positive *β* indicates selection favouring increases in the focal trait, whereas negative *β* indicates selection favouring decreases. A positive interaction term indicates selection favouring matched trait combinations (high–high and low–low), whereas a negative interaction indicates selection favouring mismatched combinations (high–low and low–high or a trade-off). Environments are defined by year (2021 and 2022) × site (Spain, Belgium, Sweden) × treatment (control *vs*. reduced precipitation). See Figure 2 and Supplementary Figure S1.

| Fruits |  |  |  |  |  |  |  |  |  |  |  |
| --- | --- | --- | --- | --- | --- | --- | --- | --- | --- | --- | --- |
| Year | Site | Treatment | $\beta$ Resistance | SE Resistance | $p$ | $\beta$ Rosette | SE Rosette | $p$ | Resistance × Rosette | SE Resistance × Rosette | $p$ |
| 2021 | Sweden | Control | 0.273 | 0.123 | 0.013 | 0.856 | 0.185 | 0.000 | 0.027 | 0.106 | 0.796 |
| 2022 | Sweden | Control | 0.060 | 0.135 | 0.650 | 0.096 | 0.139 | 0.472 | -0.011 | 0.125 | 0.928 |
| 2021 | Belgium | Control | -0.004 | 0.079 | 0.964 | 0.401 | 0.143 | 0.001 | -0.048 | 0.082 | 0.572 |
| 2022 | Belgium | Control | 0.034 | 0.079 | 0.660 | 0.859 | 0.192 | 0.000 | 0.004 | 0.081 | 0.961 |
| 2021 | Spain | Control | -0.060 | 0.076 | 0.444 | 0.792 | 0.170 | 0.000 | 0.088 | 0.089 | 0.303 |
| 2022 | Spain | Control | 0.123 | 0.093 | 0.159 | 0.341 | 0.109 | 0.000 | 0.081 | 0.084 | 0.319 |
| 2021 | Sweden | Reduced precipitation | 0.103 | 0.094 | 0.252 | 0.375 | 0.130 | 0.001 | 0.155 | 0.102 | 0.100 |
| 2022 | Sweden | Reduced precipitation | -0.278 | 0.072 | 0.001 | 0.875 | 0.199 | 0.000 | 0.151 | 0.097 | 0.093 |
| 2021 | Belgium | Reduced precipitation | 0.019 | 0.082 | 0.810 | 0.788 | 0.168 | 0.000 | -0.094 | 0.074 | 0.227 |
| 2022 | Belgium | Reduced precipitation | 0.068 | 0.076 | 0.356 | 0.551 | 0.130 | 0.000 | -0.086 | 0.064 | 0.198 |
| 2021 | Spain | Reduced precipitation | 0.042 | 0.083 | 0.602 | 0.625 | 0.148 | 0.000 | 0.040 | 0.086 | 0.636 |
| 2022 | Spain | Reduced precipitation | -0.076 | 0.100 | 0.466 | -0.455 | 0.056 | 0.000 | 0.293 | 0.168 | 0.048 |
| Stolons |  |  |  |  |  |  |  |  |  |  |  |
| 2021 | Sweden | Control | -0.167 | 0.034 | 0.000 | 0.413 | 0.062 | 0.000 | -0.032 | 0.042 | 0.444 |
| 2022 | Sweden | Control | -0.186 | 0.057 | 0.003 | 0.393 | 0.106 | 0.000 | -0.005 | 0.073 | 0.951 |
| 2021 | Belgium | Control | -0.039 | 0.033 | 0.241 | 0.504 | 0.064 | 0.000 | 0.014 | 0.035 | 0.688 |
| 2022 | Belgium | Control | 0.114 | 0.092 | 0.191 | 0.716 | 0.110 | 0.000 | -0.051 | 0.071 | 0.486 |
| 2021 | Spain | Control | 0.042 | 0.042 | 0.312 | 0.350 | 0.059 | 0.000 | 0.014 | 0.039 | 0.717 |
| 2022 | Spain | Control | -0.059 | 0.067 | 0.398 | 0.511 | 0.104 | 0.000 | 0.033 | 0.071 | 0.631 |
| 2021 | Sweden | Reduced precipitation | -0.170 | 0.033 | 0.000 | 0.180 | 0.051 | 0.000 | -0.016 | 0.035 | 0.649 |
| 2022 | Sweden | Reduced precipitation | -0.263 | 0.052 | 0.000 | 0.621 | 0.110 | 0.000 | 0.011 | 0.059 | 0.851 |
| 2021 | Belgium | Reduced precipitation | 0.030 | 0.031 | 0.322 | 0.207 | 0.043 | 0.000 | -0.048 | 0.030 | 0.122 |
| 2022 | Belgium | Reduced precipitation | 0.049 | 0.055 | 0.361 | 0.462 | 0.083 | 0.000 | -0.039 | 0.044 | 0.390 |
| 2021 | Spain | Reduced precipitation | 0.026 | 0.040 | 0.504 | 0.254 | 0.055 | 0.000 | 0.016 | 0.038 | 0.673 |
| 2022 | Spain | Reduced precipitation | -0.214 | 0.053 | 0.000 | 0.129 | 0.079 | 0.084 | 0.134 | 0.069 | 0.039 |

**Supplementary Figure S1.**
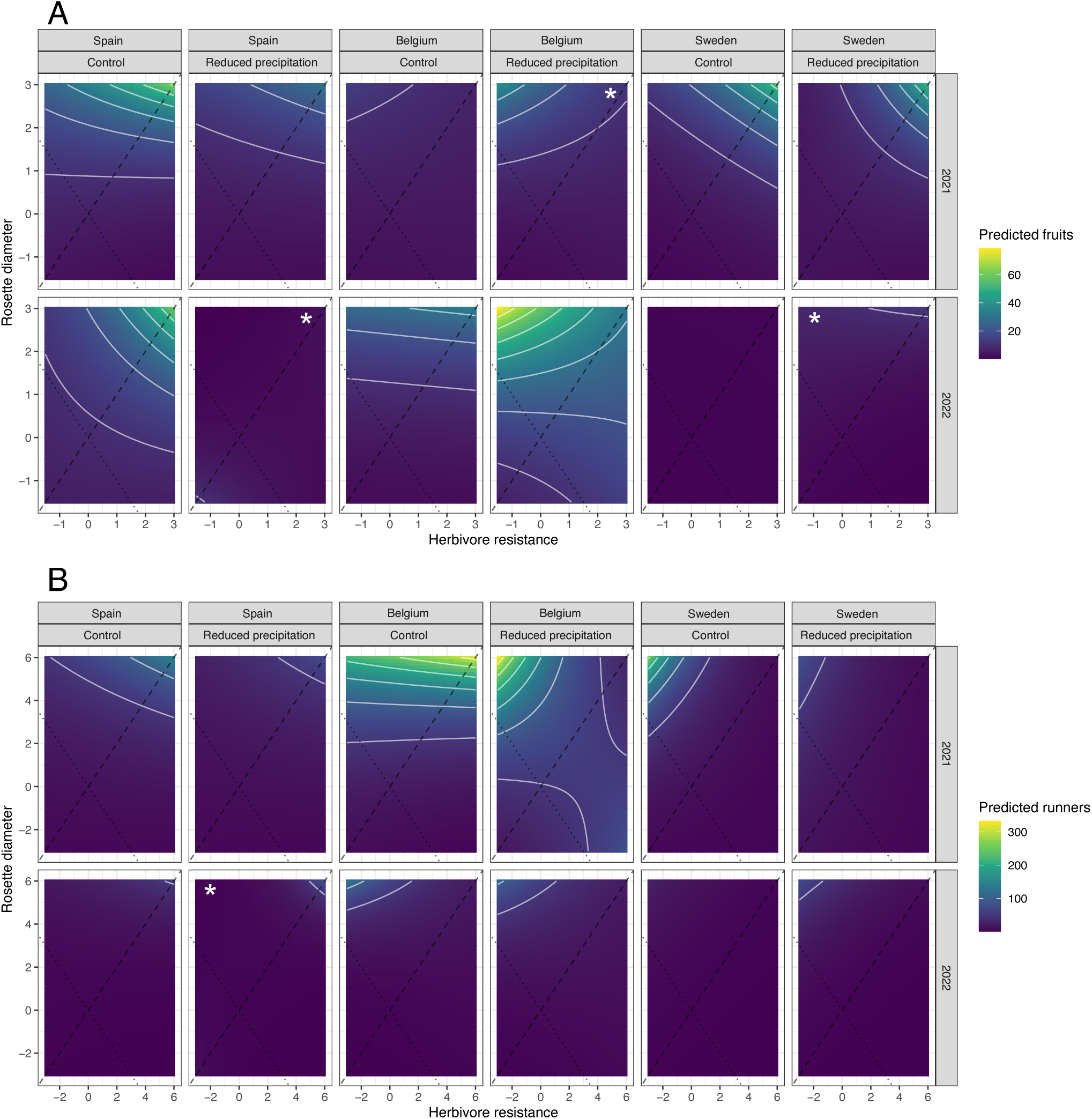
Correlational selection surfaces between rosette diameter and herbivore resistance across environments; sites (Spain, Belgium and Sweden), treatments (control and reduced precipitation) and years (2021 and 2022). (A) Predicted fruit production and (B) predicted runner production are shown as functions of herbivore resistance (x-axis) and rosette diameter (y-axis). For each environment (site × treatment × year), the heatmap shows predicted fitness (lighter colours indicate higher values; see colour bars), and contour lines denote iso-fitness contours. Dashed and dotted diagonals indicate reference axes representing contrasting trait combinations (high–high *vs*. high–low). White asterisks depict significance of the correlational selection. See Supplementary Table S3.

**Supplementary Table S4.**
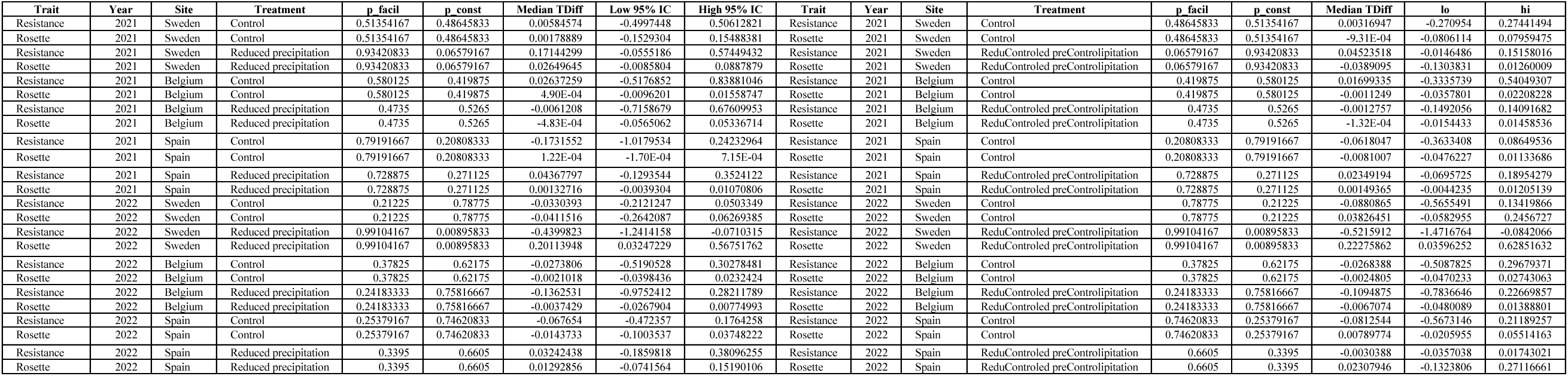
Genetic facilitations and constraints across environments (i.e., every site × treatment × year combination). Genetic covariances were classified as facilitating evolutionary change for resistance or rosette diameter when FC > 0, indicating that covariances increase the magnitude of the predicted response in the direction of selection. This is indicated by the *p_facil* column, which is showing the posterior probability of *direction-aware facilitation* (FC > 0). Conversely, covariances were classified as constraining when FC < 0 or p_const showing the posterior probability of *direction-aware constraint* (FC < 0). Posterior median of Tdiff is showing the *effect size* or how much covariances change the predicted response of that trait. 2.5% and 97.5% quantiles of TDiff (a 95% credible interval for TDiff). Significant facilitation: p_facil ≥ 0.90, significant constraint: p_const ≥ 0.90. See also methods and Figure 4.

